# Non-invasive imaging of salicylic and jasmonic acid activities *in planta*

**DOI:** 10.1101/2025.02.17.636591

**Authors:** Anastasia V Balakireva, Tatiana A Karataeva, Michael Karampelias, Tatiana Yu Mitiouchkina, Jan Macháček, Ekaterina S Shakhova, Maxim M Perfilov, Olga A Belozerova, Kseniia A Palkina, Nikola Drazna, Zuzana Vondrakova, Aubin Fleiss, Liliia I Fakhranurova, Nadezhda M Markina, Viktor V Morozov, Evgenia N Bugaeva, Galina M Delnova, Vladimir V Choob, Ilia V Yampolsky, Jan Petrášek, Alexander S Mishin, Karen S Sarkisyan

## Abstract

Jasmonic and salicylic acids are the major hormones involved in plant response to pests and pathogens. Here, we engineered autoluminescent plants that report activity of these hormones with up to 53-fold contrast. Using consumer-grade cameras, we imaged reporter *Arabidopsis thaliana* and *Nicotiana benthamiana* plants throughout normal development, and in response to attacks of pests and pathogens.

---

Salicylic acid (SA) and jasmonic acid (JA) are plant hormones involved in regulating plant defence mechanisms and development. Salicylic acid primarily mediates response to biotrophic pathogens ^1^, while jasmonic acid orchestrates defence responses against necrotrophic pathogens and herbivores ^2^. Both hormones play essential roles in modulating stress responses, growth, and development. The relationship between salicylic and jasmonic acid pathways is mostly described as antagonistic ^3^. This antagonism can be exploited by pathogens. For example, *Pseudomonas syringae* suppresses salicylic acid response by elevating the levels of jasmonic acid ^4^, while whiteflies tilt the response towards salicylic acid ^5^. Interplay of jasmonic and salicylic acids regulates local and systemic responses to pest attacks; these interactions are in turn affected by other hormones, including ethylene and auxin ^6,7^.

How active are these hormones in different plant tissues in various ecological contexts, how local is their action, and how does it develop over time? An non-invasive imaging approach offering high spatial and temporal resolution could enable studies aimed at answering these questions. Existing approaches are generally limited to the use of fluorescent probes that require external illumination ^8^, transcriptional luminescent reporters coupled with exogenous addition of luciferins ^9^, or mass-spectrometry-based techniques that are typically invasive and rely on expensive equipment ^10^.

Autoluminescence pathways produce luminescence substrate in the studied organism throughout its lifespan ^11–18^, potentially allowing non-invasive imaging of molecular physiology at the organism level. In the fungal bioluminescence pathway (**Figure 1a**), caffeic acid, a molecule present in all land plants, is converted into luciferin ^19–21^. In plants, the pathway can be reconstituted by overexpressing five fungal genes: hispidin synthase HispS, a phosphopantetheinyl transferase such as **NpgA** from *Aspergillus nidulans*, hispidin-3-hydroxylase H3H, luciferase **Luz**, and caffeylpyruvate hydrolase **CPH** ^13,16,22–24 13,25^. Here, we adapted these genes to create transcription-based autoluminescence reporters that visualise hormone activities in transgenic plants.

**Figure 1.**
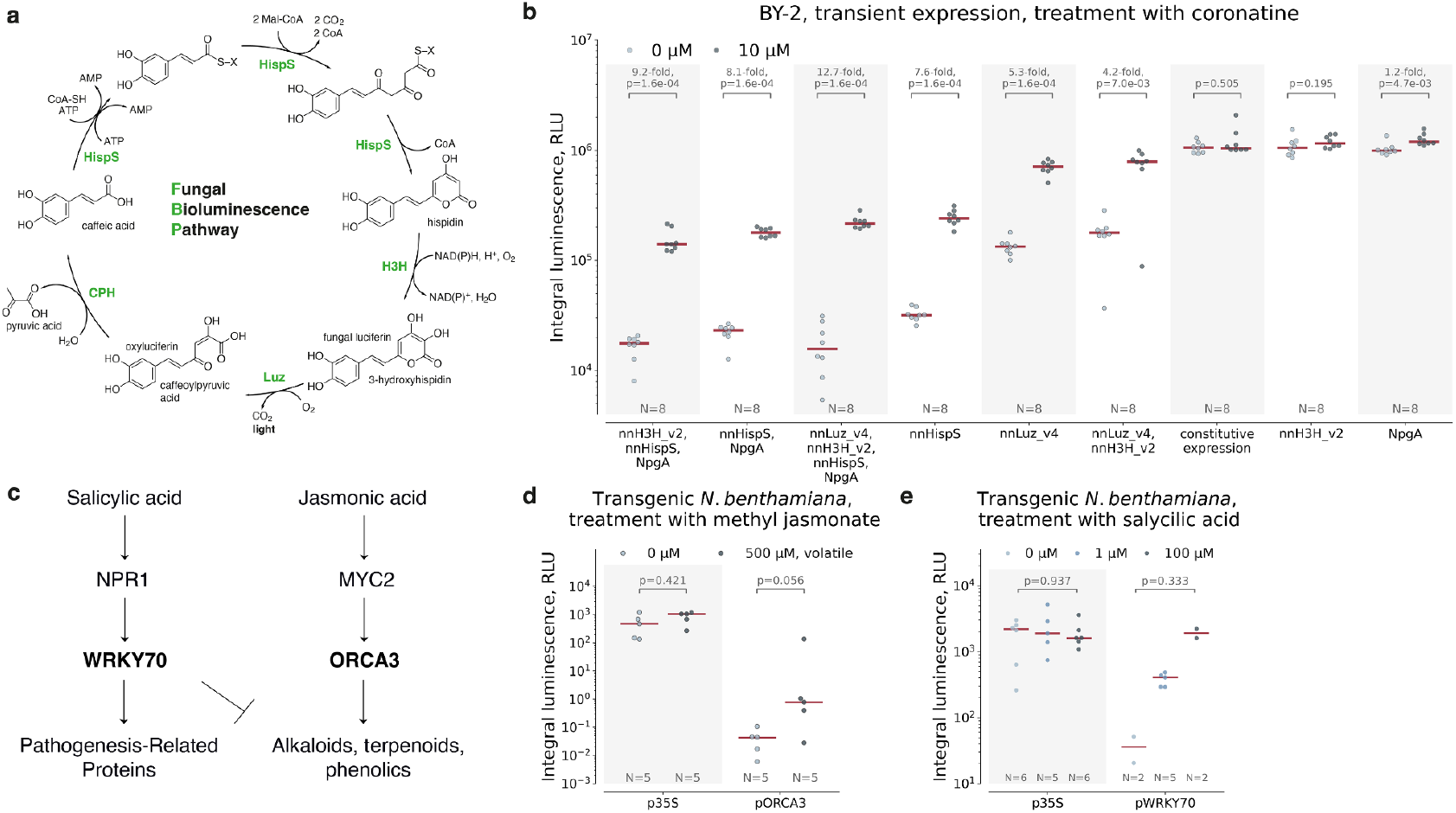
Prototyping the design of reporters based on the fungal bioluminescence pathway. **a.**Biochemical reactions of the fungal bioluminescence pathway are catalysed by hispidin synthase **HispS**, hispidin-3-hydroxylase **H3H**, luciferase Luz and putative caffeoyl pyruvate hydrolase **CPH**. For its activity, **HispS** requires a post-translational modification by a phosphopantetheinyl transferase, such as **NpgA** from *Aspergillus nidulans*. **b**. Performance of various architectures of transcriptional reporters in cell packs from BY-2 cells. Genes controlled by the jasmonate-responsive promoter pORCA3 are indicated on the horizontal axis. Concentrations of jasmonic acid agonist coronatine, that was used as an inducer, are specified in the legend. **c**. Simplified presentation of signalling pathways induced by salicylic and jasmonic acids. WRKY70 promotes salicylic acid response, while inhibits jasmonic acid signalling. ORCA3 is a transcription factor which expression is activated by jasmonates. The ORCA3 promoter contains a jasmonate-responsive element that operates through interactions with transcription factors like AtMYC2. Inducibility of pORCA3-nnLuz (**d**) and pWRKY70-nnLuz (**e**) in transgenic *Nicotiana benthamiana*. The red line is the median, the coloured points represent individual data points. The difference between mean values and *p*-values of post-hoc two-sided Mann–Whitney U tests are indicated above the brackets between the box plots. N = 2-8 biologically independent samples.

We first tested what genetic architecture would produce optimal light emission without compromising the dynamic range. Previous studies pointed to HispS-catalysed reaction as the bottleneck of the pathway^23,24,26^, suggesting that inducible expression of hispidin synthase could provide tighter control over luminescence. However, due to weak activity of hormone-sensitive promoters, insufficient levels of hispidin synthase might result in low signal-to-noise ratio.

We tested different designs encoding bioluminescence pathway, using inducible promoters previously validated in our laboratory: jasmonic-acid-sensitive *Catharanthus roseus* pORCA3 [ref ^27,28^] and cytokinin-sensitive promoter *Arabidopsis thaliana* pARR6 [ref ^29^]. In these designs, one or more genes of the pathway were controlled by pORCA3 or pARR6, while the remaining genes were expressed under strong constitutive promoters. In plant cell packs made of *Nicotiana tabacum* BY-2 cells ^30^, luminescence increased in most pathway designs upon induction (**Figure 1b, Supplementary Figure 1**). The HispS-NpgA-inducible design indeed showed highest activation and lowest background signal, in agreement with the previously reported data ^23,24,26^. However, its luminescence in the induced state was dim, while Luz-inducible design demonstrated brighter luminescence, showing a comparable level of activation.

We thus settled on the design where only the luciferase was driven by a hormone-sensitive promoter, while the rest of the genes were expressed constitutively. To construct a jasmonate reporter, we placed nnLuz_v4 gene ^23^ under control of the promoter and terminator of *Catharanthus roseus* ORCA3 gene ^27^. For salicylic acid, we used the corresponding regulatory elements from *Arabidopsis thaliana* WRKY70 gene [ref ^29^] activated by salicylic acid and repressed by jasmonates ^31^. pORCA3-inducible reporter responded to treatment with methyl jasmonate with an increase in luminescence (**Supplementary Figure 3**). In the case of pWRKY70, luminescence of untreated samples was high, likely reflecting upregulation of WRKY70 following agrobacterial infection in our assay ^32^ (**Supplementary Figure 3**).

We continued evaluating the reporters in transgenic *Nicotiana benthamiana* and *Arabidopsis thaliana* lines. We first created luciferin-producing master lines **NB237** and **AT8462**, respectively, that constitutively expressed all bioluminescence genes, except luciferase (**Supplementary Table 2, Supplementary Figure 4**). These lines were used to create plants with pORCA3-, pWRKY70- or p35S-driven expression of luciferase. *N. benthamiana* plants expressing pWRKY70-inducible luciferase responded to treatment with salicylic acid in a concentration-dependent manner, showing up to 53-fold overall increase in light emission (**Figure 1e**, the setup is shown at **Supplementary Figure 5**). Accordingly, pORCA3 reporter plants showed an increase in luminescence upon treatment with methyl jasmonate (**Figure 1d**).

We then tested these plant lines in wounding experiments and upon pathogen attack. Wounding and infection with necrotrophic pathogens is known to raise the level of jasmonic acid. Infection with biotrophic pathogens, in turn, leads to accumulation of salicylic acid ^33^. In accordance with this, wounding of JA-reporting *N. benthamiana* plants induced luminescence locally (**Supplementary Figures 7 and 9, Supplementary videos 1 and 2**). Quick local induction of luminescence at infiltration sites was also observed in lines constitutively expressing the luciferase, likely reflecting activation of phenylpropanoid metabolism ^16^. Analysis of leaves with LC-MS confirmed an increase in levels of jasmonic acid characteristic to the response to wounding (**Supplementary Figure 6**).

In turn, SA-reporting plants reacted to infiltration with *Agrobacterium tumefaciens* but not with the buffer (**Supplementary Figure 8, Supplementary video 3**). Upon infiltration with hemibiotrophic pathogen *Pseudomonas savastanoi*, JA-reporting plants responded primarily to wounding itself (**Supplementary Figure 9a**). In contrast, luminescence of SA reporters increased ∼15-fold specifically upon infiltration with the pathogen (**Supplementary Figure 8a**), in accordance with salicylic acid response described in the literature ^34,35^. In the case of a necrotrophic bacterium *Pectobacterium carotovorum* we observed a striking increase in luminescence emitted from the vasculature of non-infiltrated, or systemic, leaves of JA-reporting plants (**Figure 2a, Supplementary Video 1, Supplementary Figure 9a**), also in accordance with existing data ^2^. This vasculature signal likely visualises early stages of induced systemic resistance.

**Figure 2.**
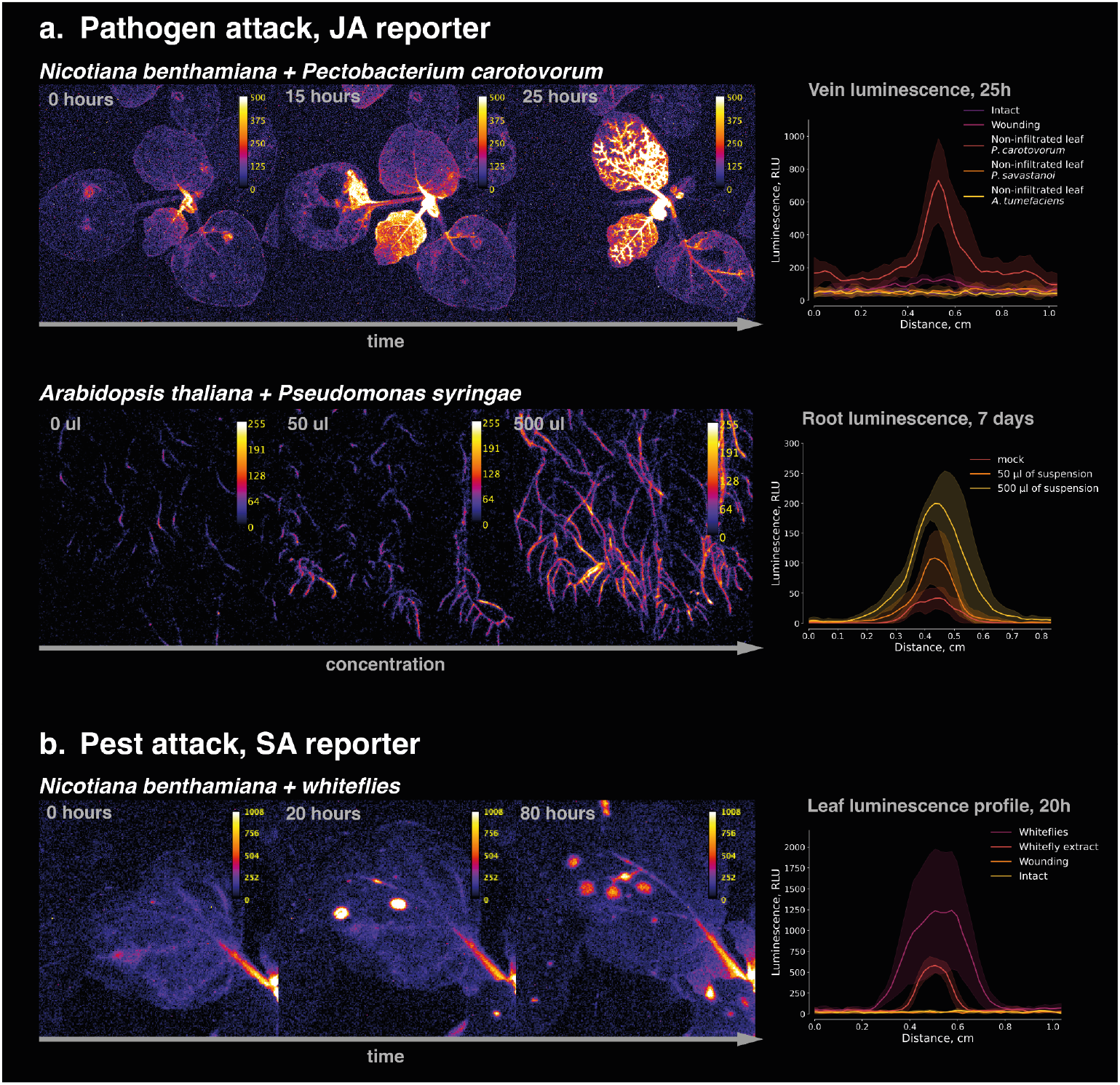
Performance of reporters of salicylic and jasmonic acids in transgenic *Nicotiana benthamiana* or *Arabidopsis thaliana* plants. **a.** Luminescent imaging of JA-reporting N. benthamiana (up) or A. thaliana (bottom) upon infection with *Pectobacterium carotovorum* or *Pseudomonas syringae* pv. tomato DC3000, respectively. Pst50 and Pst500 – 50 μl or 500 μl of suspension in 50 mL of buffer, respectively. Plots indicate *N. benthamiana* non-infiltrated leaf veins brightness compared to tissues nearby at 25 hours post infection (up, N = 6) or *A. thaliana* root brightness treated with different concentrations of bacteria, 7 days post treatment (bottom, N = 6-7 roots). **b**. Luminescent imaging of SA-reporting plants infested with whiteflies. Plots indicate the brightness of leaves’ local responses to whitefly bites. N = 3-7 local responses.

A similar behaviour was observed in *Arabidopsis thaliana* reporter lines. We incubated plants with *E*.*coli, A*.*tumefaciens*, and *P. syringae pv. tomato DC3000*, tracking their luminescence for 7 days. While JA-reporting plants showed no increase in luminescence upon treatment with *E*.*coli* or *A*.*tumefaciens*, they demonstrated dose-dependent response to *P. syringae* in the infected lateral roots (**Figure 2a, Supplementary Figure 10**). The induction of the JA pathway is expected since *P*.*syringae* produces coronatine—jasmonate mimic—to exploit the interplay between JA and SA pathways ^36^.

SA-reporting *Arabidopsis* plants did not respond to pathogenic *P. syringae*, showed minor response in the roots to high concentrations of *E*.*coli*, and responded to *A*.*tumefaciens*. During the first days of infection, *A*.*tumefaciens* reduced luminescence in the roots, and induced it in the shoots. After the first 4 days, pathogenesis advanced, resulting in halted development and chlorotic leaves (**Supplementary Figure 11**).

Taking advantage of the abundance of greenhouse whiteflies *Trialeurodes vaporariorum* in one of our facilities, we also aimed to monitor response of *N*.*benthamiana* plants to their attack in a 7-day experiment (**Figure 2a, Supplementary Figures 7-9, 12-13, Supplementary Videos 4-12**). Previous studies showed that whitefly salivary protein effector Bt56 activates salicylic acid signalling, lowering the level of jasmonic acid and restraining plant response against the insect. We released whiteflies onto reporter plants and imaged them for 7 days (**Supplementary Videos 4-12**). No luminescence response was observed from jasmonate-reporting plants (**Supplementary Figure 9b**). In contrast, salicylic acid reporter plants developed light-emitting spots, likely visualising local response to whitefly bites (**Figure 2b**). When we wounded leaves with needles dipped in the control buffer, we did not observe a response. But if the needles were dipped in homogenised whiteflies, wounding led to a notable response (**Supplementary Figure 8b**), indicating specificity to whiteflies. On the sixth day of imaging, we registered an overall increase in brightness of plants bitten by whiteflies, potentially illustrating the development of a systemic acquired resistance triggered by the Bt56 effector ^37,38^ (**Supplementary Figure 13**). LC-MS analysis confirmed an increase of salicylic acid concentration in whitefly-bitten plants (**Supplementary Figure 14**).

We then performed long-term imaging of hormonal activities in *N*.*benthamiana* upon normal development. We followed plant development for over 7 weeks, starting from 4-leaf plants (**Supplementary Videos 13-15**). The JA-sensitive line was dim throughout the imaging experiment and became significantly brighter at flowering (**Supplementary Figure 15**), demonstrating bright luminescence from floral primordia and the tube of the mature flower, whereas in limb and sepals the brightness rapidly decreases after the flower opening, in agreement with existing data on elevated jasmonate activity in reproductive tissues ^39^. Leaf veins of these plants emitted little light (**Supplementary Figure 15**). In contrast, SA-sensitive plants showed brighter luminescence from leaf vasculature and almost no light from flowers (**Supplementary Figure 15**) ^40^. The contrasting pattern of luminescence from leaf vasculature in two reporter lines may reflect the antagonistic relationship between salicylic acid and jasmonates. Control plants constitutively expressing bioluminescence genes (**Supplementary Figure 15**) had evenly distributed luminescence emitted by leaves, flowers and stems.

Although approaches based on autoluminescence have drawbacks, such as dependence of signal on metabolic activity of a tissue, they can provide insightful observations that may be further tested by orthogonal approaches. Ability to perform luminescence imaging without exogenous substrate and with inexpensive off-the-shelf consumer cameras enables studies of plant molecular physiology in the field, at scale, and in low-resource settings. We expect reporters described here to be useful for visualising effects of pest and pathogen pressure in the natural ecological environment, and to contribute to deeper understanding of plant physiology.

## Acknowledgements

This study was partially funded by Planta (planta.bio) and Light Bio (light.bio). We thank Milaboratory (milaboratory.com) for the access to computing and storage infrastructure. The Synthetic Biology Group is funded by the MRC Laboratory of Medical Sciences (UKRI MC-A658-5QEA0). This work was supported by UKRI Biotechnology and Biological Sciences Research Council through the International Science Partnerships Fund (ISPF) [grant number UKRI249]. Whole-plant imaging experiments were funded by RSF, project number 24-74-10105 (https://rscf.ru/project/24-74-10105/). Plant transformations were funded by RSF, project number 24-74-10087 (https://rscf.ru/project/24-74-10087/). Works in Arabidopsis were supported by the CSF project n. 21-20936J (ZV, MK, JP), Czech BioImaging project n. LM 2023050, and the HORIZON-WIDERA-2022-TALENTS-02 project n. 101090293 (MK).

## Author Contributions

Performed experiments: TAK, ESS, KAP, TYM, KM, MVV, MK, NMM, LIF, GD, OAB, ENB.

Planned experiments: AVB, TAK, ESS, KAP, TYM, MK, KSS, ASM, VVC.

Analysed data: MMP, AVB, KSS, ASM, VVC.

Proposed and directed the study: KSS, JP, ASM, IVY.

Reviewed the paper draft: all authors.

## Materials and methods

### Design and assembly of genetic constructs

Coding sequences of bioluminescence genes were optimised for the expression in *Nicotiana benthamiana* and ordered synthetically (**Supplementary Table 1**). Golden Gate assembly was performed in the T4 ligase buffer (Thermo Fisher) containing 10 U of T4 ligase, 20 U of either BsaI or BpiI (Thermo Fisher) and ∼100 ng of each DNA part. Typically, Golden Gate reactions were performed according to ‘troubleshooting’ cycling conditions described in [ref ^41^]: 25 cycles (90 s at 37 °C, 180 s at 16 °C), then 5 min at 50 °C and 10 min at 80 °C.

Correct DNA assembly was typically confirmed by Sanger sequencing, and in some cases additionally by Nanopore or Illumina-based whole plasmid sequencing. DNA assembly and whole-plasmid sequencing was typically ordered from Cloning Facility (cloning.tech).

To generate stable *Nicotiana benthamiana* plants expressing either salicylic- or jasmonic-acid-responsive reporters, we first developed a “luciferase-less” masterline carrying genes necessary for fungal luminescence, except the luciferase gene. To obtain such masterline we assembled Level P plasmid pNK091 carrying (1) nnCPH gene under the control of 0.4 kb constitutive 35S promoter from cauliflower mosaic virus with 5′ untranslated region of TMV omega virus and ocs terminator from *Agrobacterium tumefaciens*; (2) wild-type version of nnH3H gene under the control of constitutive FMV promoter from figwort mosaic virus and nopaline synthase terminator from *Agrobacterium tumefaciens*; (3) nnHispS gene under the control of 0.4 kb constitutive 35S promoter from cauliflower mosaic virus with 5′ untranslated region of TMV omega virus and ocs terminator from *Agrobacterium tumefaciens*; (4) NpgA gene under the control of the constitutive CmYLCV 9.11 promoter from Cestrum yellow leaf curling virus and ATPase terminator from *Solanum lycopersicum*; (5) kanamycin resistance cassette driven by pNos promoter and ocs terminator from *Agrobacterium tumefaciens*.

The master line **NB237** was then further transformed with plasmids encoding phytohormone-sensitive luciferase-carrying transcription units. To obtain phytohormone-sensitive plants masterline was transformed with the Level M plasmid carrying hygromycin resistance cassette driven by pNos promoter and ocs terminator from *Agrobacterium tumefaciens* and nnLuz_v4 ^23^ under the control of WRKY70 promoter and terminator in the case of salicylic acid reporter (plasmid ID: M7691) or under the control of ORCA3 promoter or terminator – in the case of jasmonic acid reporter (plasmid ID: M7138). In control samples nnLuz_v4 gene was placed under the constitutive 0.4 kbp 35S promoter and AtAct2 terminator (plasmid ID: pNK3558).

### Transformation of *Agrobacterium tumefaciens*

Plasmids were transformed into competent cells of *Agrobacterium tumefaciens* AGL0 ^42^, and clones were selected on LB (Luria-Bertani) agar plates containing 50 mg/L of rifampicin and an additional antibiotic, depending on the plasmid used for transformation (200 mg l−1 of carbenicillin, 50 mg ml−1 of kanamycin or 100 mg ml−1 spectinomycin). Individual colonies were then inoculated into 10 ml of LB medium containing the same concentration of antibiotics. After overnight incubation at 28 °C with shaking at 220 rpm, cultures were centrifuged at 2,900g, resuspended in 25% glycerol and stored as glycerol stocks at −80 °C.

### Growth conditions for *Arabidopsis thaliana*

Seeds were surface sterilised with 70% ethanol for 15 minutes and washed twice with 100% ethanol. Seeds were dried in a laminar flow and then sown on the surface of solid (10% plant agar) half-strength MS medium with 1% sucrose. Sown seeds were vernalized for 2 days at 4°C and then transferred to grow vertically at 22°C.

### Validation of reporters in plant cell packs based on *Nicotiana tabacum* BY-2 cell culture

BY-2 cell culture was grown in BY-2 medium (Murashige and Skoog (MS) with 0.2 mg l−1 2,4-dichlorophenoxyacetic acid, 200 mg l−1 KH2PO4, 1 mg l−1 thiamine, 100 mg l−1 myo-inositol and 30 g l−1 sucrose) at 27 °C by shaking at 130 rpm in darkness, with 2 ml of 1-week-old culture being transferred into new 200 ml of BY-2 medium every week ^43^.

Transformations of BY-2 cell packs were made according to a protocol adapted from ^44^. One-week-old BY-2 culture was pelleted in black 96-well plates to create cell packs that were infiltrated by a mixture of several agrobacterial strains containing binary vectors. One of the strains encoded silencing inhibitor P19 (OD_600_0.2), and other encoded bioluminescence genes and a reporter (ARR6-, WRKY70-, ORCA3-nnLuz_v4 variants) gene (OD_600_0.5). Comparison of reporters was done by co-infiltrating BY-2 cell packs with agrobacteria individually encoding bioluminescence enzymes.

Salicylic acid and jasmonic acid reporter functionality was tested in BY-2 cell packs. In the case of pWRKY70-nnLuz_v4-WRKY70_T, the cell packs were treated with 100 μM salicylic acid solution. The 100 μl of salicylic acid solution was applied on the cell packs, the excess of the solution was removed with centrifugation for 1 min, 500g. In the case of pORCA3-nnLuz_v4-ORCA3_T, the cell packs were treated with methyl jasmonate emitted from a surface of 500 μM solution placed in between the rows of 96-well plates.

After the treatments the plates were incubated at 80% humidity at 22 °C and immediately imaged for 48 hours using Sony Alpha ILCE-7M3 camera and 35-mm T1.5 ED AS UMC VDSLR lens (Samyang, ∼f/1.4) with an exposure of 5–30 s and ISO 400, 3,200, and 20,000.

Processing of images was performed using custom Python scripts (Python version 3.10.12). For luminescence quantification values in the wells were used.

### Agrobacterium-mediated transformation of *Nicotiana benthamiana*

*Agrobacterium tumefaciens* strains AGL0 carrying plasmid pNK091 (for creation of a masterline) or plasmids M7138, M7691, pNK3558 (for the development of reporter lines) were grown in flasks on a shaker overnight at 28 °C in LB medium supplemented with 25 mg l−1 rifampicin and 50 mg l−1 kanamycin or hygromycin, respectively. Bacterial cultures were diluted in liquid MS medium to an optical density of 0.6 at 600 nm. Leaf explants used for transformation experiments were cut from 2-week-old *Nicotiana benthamiana* plants and incubated with bacterial culture for 20 min. Leaf explants were then placed onto filter paper overlaid on MS medium (MS salts, MS vitamins, 30 g l−1 sucrose and 8 g l−1 agar, pH 5.8) supplemented with 1 mg l−1 6-benzylaminopurine and 0.1 mg l−1 indolyl acetic acid. Two days after inoculation, explants were transferred to the same medium supplemented with 500 mg l−1 cefotaxime and 75 mg l−1 kanamycin or hygromycin. Regeneration shoots were cut and grown on MS medium with antibiotics.

### Transformation of *Arabidopsis thaliana*

Arabidopsis thaliana (ecotype Columbia 0) was transformed with floral dip ^45^ with a binary vector carrying the transcriptional units for the ectopic production of the fungal luciferin, and the recycling of the luciferin’s oxidization product (caffeoylpyruvic acid). After 4 generations of antibiotic resistance (kanamycin), homozygous plants were transformed with a binary vector with a transcriptional unit regulation of the expression of the fungal luciferase by the ORCA3 or WRKY70 promoters and corresponding terminators. Transgenic plants were selected for resistance to hygromycin, and bioluminescence, captured by the Sapphire (Azzure) bioimager (exposure time was 2 minutes). Second-generation, homozygous plants were used further for experiments.

### Validation of phytohormone reporters in transgenic *Nicotiana benthamiana*

All experiments were carried out on T1 plants at two developmental stages: 1-week-old seedlings grown *in vitro* in Petri dishes, and 3-4-week-old plants grown in peat tablets. In all experiments 3 stable lines of *Nicotiana benthamiana* were used: they carried transcription units of fungal bioluminescence genes: pNos-KanR-ocsT | p35S-nnHispS-ocsT | pCmYLCV-npgA-ATPT | p35S-nnCPH-ocsT | pFMV-nnH3H_wt-nosT – and either salicylic acid reporter pWRKY70-nnLuz_v4-WRKY70_T, jasmonic acid reporter pORCA3-nnLuz_v4-ORCA3_T, or control p35S_0.4kbp-nnLuz_v4-act2. For each experiment, we used 3-5 plants as replicates.

### Wounding

To induce jasmonic acid signalling we damaged the stems of plants grown in vitro or the leaves of the plants by cutting them with scissors.

### Salicylic acid and methyl jasmonate treatment

To induce salicylic acid signalling we sprayed the plants with 100 μM salicylic acid solution. To induce jasmonate signalling we treated the plants with methyl jasmonate emitted from a surface of a cotton wool soaked in 5 mM solution and placed next to the plants.

### Infection with bacteria

We infected transgenic plants with bacteria using different infection strategies. For the experiments we used a hemibiotrophic pathogen *Pseudomonas savastanoi* (a pathovar of *Pseudomonas syringae)*, and necrotrophic bacteria *Pectobacterium carotovorum*. To infect the plants we first damaged the stems and then applied the pathogens in wounded areas. In parallel, the effect of the agrobacterium *Agrobacterium tumefaciens* strain AGL0 infection was studied. A 10 mM MgSO_4_, 0.01% Silwet L-77 solution in which bacteria were resuspended was used as a control. The OD_600_ of bacteria was 0.2.

### Whitefly infestation

The greenhouse whitefly *Trialeurodes vaporariorum* was used for the experiments. A total of 100 whiteflies per plant was applied on assayed *Nicotiana* lines. These plants were covered with a glass fish tank to trap the insects. To imitate the whitefly bite we pricked the leaves with sterile needles dipped in either 0.1 M phosphate buffer pH 7.0 (mock) or the whitefly extract. The extract was prepared by homogenising 50 whiteflies in 2 mL of the 0.1 M phosphate buffer pH 7.0.

### Luminescent imaging

The plants were imaged for 48 hours using Sony Alpha ILCE-7M3 camera and 35-mm T1.5 ED AS UMC VDSLR lens (Samyang, ∼f/1.4) with an exposure 30 s and ISO 400, 3,200, and 20,000. We typically imaged plants every 30 minutes.

### Long-term timelapse luminescent imaging of phytohormone-sensitive plants

Long-term timelapse luminescent imaging of hormone-sensitive plants was performed for 1.5 month using Sony Alpha ILCE-7M3 camera and 35-mm T1.5 ED AS UMC VDSLR lens (Samyang, ∼f/1.4) with an exposure of 5–30 s and ISO 400, 3,200, and 20,000. The snapshots were captured every 30 minutes.

Processing of images was performed using FiJi ImageJ distribution (version 1.53t) and custom Python scripts. For luminescence quantification mean values in the region of interest after background subtraction were used.

Background subtraction was performed using the following formula: signal = signal_raw_ − background_mean_.

### Validation of phytohormone reporters in transgenic *Arabidopsis thaliana*

The bacteria used in this study were *Escherichia coli* (XL1 blue), *Agrobacterium tumefaciens* (GV2260), and *Pseudomonas syringae* pv. tomato DC3000. Single colonies were grown on solid LB medium inoculated 1.5 ml of liquid LB and the cultures were incubated in a shaker at 28 or 37°C, for *A. tumefaciens* and *P. syringae*, or *E*.*coli*. After 20 hours of incubation, bacterial cultures were centrifuged at 1500g for 5 minutes, LB was discarded and pelleted bacteria were resuspended in autoclaved distilled water to O.D. 1. Infection plates were set with 50 ml of solid plant growth medium inoculated with 50 or 500μl of bacterial suspensions while warm and liquid. Six-day-old plants grown vertically were transferred to plates with medium inoculated as mentioned above. One plate did not have bacteria and served as a control.

### Imaging of *Arabidopsis thaliana*

Plants grown on media with bacteria were photographed in the indicated days (d.a.t. days after transfer) with a Samyang 35mm f/1.4 AS UMC lens attached to a Nikon Z6 ii camera body. Imaging included a photo in a well-lit condition (f/1.4, ISO 200, 1/160 seconds) and a photo in complete darkness (f/1.4, ISO 51200, 60 seconds). Plates with plants were placed horizontally for less than 2 minutes while imaging. Image processing by ImageJ software includes isolation of the green channel, denoising by outliers removal, and intensity increase.

### Data presentation and statistical analysis

Most of the data are plotted as medians and coloured individual data points using Seaborn (https://seaborn.pydata.org/, ver. 0.13.2) and Matplotlib (https://matplotlib.org/, ver. 3.8.0) packages, using Python version 3.10.12. Pairwise post-hoc two-sided Mann–Whitney U tests (Scikit-posthocs package ^46^, version 0.10.0) were computed. Sample numbers (N) are reported in the figures.

### LCMS experiments

### Chemicals

The analytical standards of jasmonic acid, salicylic acid were purchased from Sigma-Aldrich® (≥98.0), hispidin was chemically synthesised and tested for purity in house (>95.0). Standard solution of two components was prepared in a 50% acetonitrile. HPLC-grade acetonitrile was purchased from J.T.Baker. Deionized water was obtained from a Milli-Q System (USA) and acetic acid was purchased from Sigma-Aldrich® (≥98.0).

### Sample preparation

Fresh leaves were homogenised in liquid nitrogen and lyophilised. Extracts were obtained from 25 mg of dry plant biomass using 1 mL of 70% methanol, filtered through 0.45 μm GF/PVDF (Phenex) filter and lyophilised in miVac machine. Dry residues were reconstituted in 100 µL of 70% methanol by vortexing, and transferred for LC-MS analysis.

### LC-MS analysis

LC-MS analysis was carried out on an Ultimate 3000 RSLC HPLC system connected to a QExactive Plus mass spectrometer (Thermo Fisher Scientific, USA). Samples were separated on a Gemini C18 3 μm NX LC column 100*2.1 mm (Phenomenex) at 200 ul/min flow rate. Separation was done by a linear gradient of 90% acetonitrile in water, 10 mM ammonium formate, 0.1% formic acid (Buffer B) in 99.9% H_2_O, 10 mM ammonium formate, 0.1% formic acid (Buffer A): 1% B at 0 min, 50% B at 3 min, 99% B at 8 min, followed by 3 min wash at 99% B and 2 min equilibration at 1% B before the next run. UV data was collected at 220 nm. MS1 and MS2 spectra were recorded at 30K and 15K resolution respectively with HCD fragmentation. Raw data were collected and processed on Thermo Xcalibur Qual and Skyline software. The MS peaks were extracted at a mass tolerance of 5 ppm. For compound quantification the corresponding peak area for each sample was used.

