## Supplementary materials for "Non-invasive imaging of salicylic and jasmonic acid activities *in planta*"

### Supplementary Information

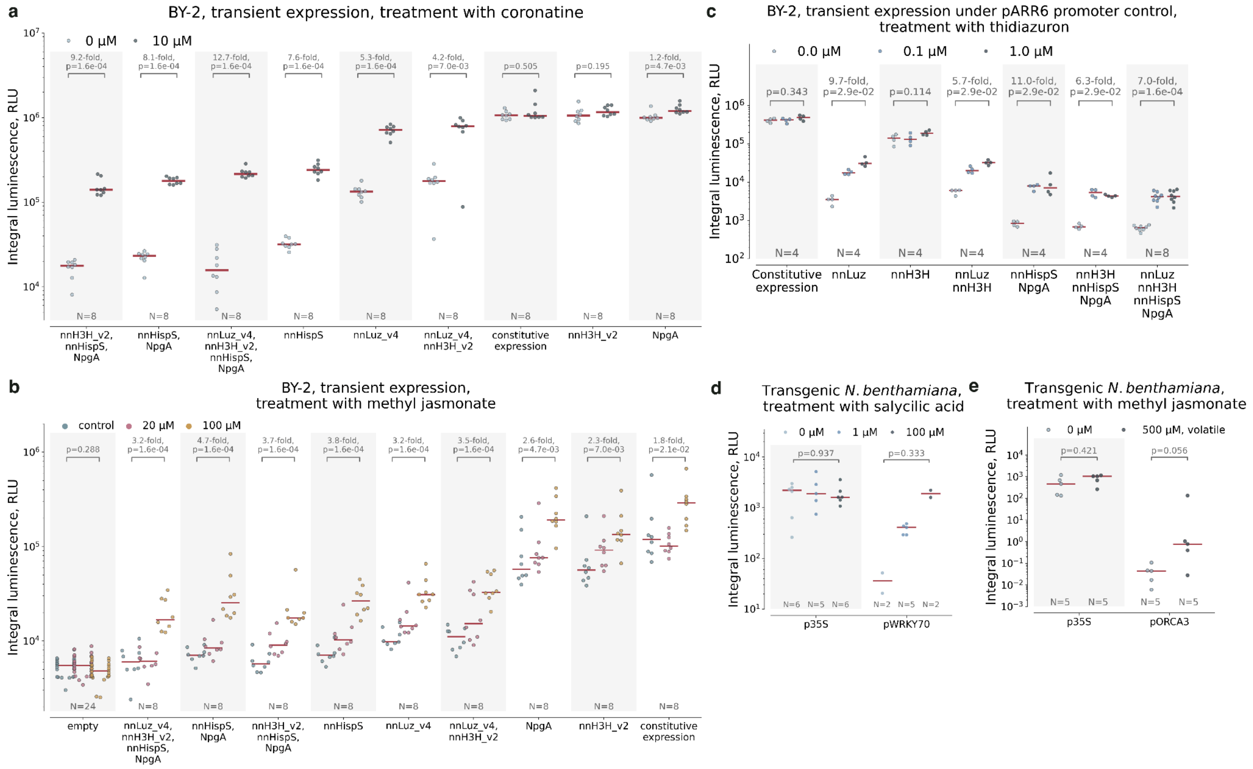


**Supplementary Figure 1.** Selection of a fungal bioluminescence gene as a reporter of phytohormone activity upon treatment with the inducer in BY-2 cell packs.

**a.** Performance of various architectures of transcriptional reporters in cell packs from BY-2 cells. Genes controlled by the jasmonate-responsive promoter pORCA3 are indicated on the horizontal axis. Concentrations of jasmonic acid agonist coronatine, that was used as an inducer, are specified in the legend. **b**. Treatment of BY-2 cell packs expressing fungal bioluminescence genes put under pORCA3 with volatile methyl jasmonate. The red line is the median, the coloured points represent individual data points. The difference between mean values and *p*-values of post-hoc two-sided Mann–Whitney U tests are indicated above the brackets between the box plots. N = 8-24 plant cell packs. **c**. Treatment of BY-2 cell packs expressing fungal bioluminescence genes put under pARR6 with 0.1 µM and 1 µM thidiazuron solution. The red line is the median, the coloured points represent individual data points. The difference between mean values and *p*-values of post-hoc two-sided Mann–Whitney U tests are indicated above the brackets between the box plots. N = 4-8 plant cell packs. Inducibility of pWRKY70-nnLuz (**d**) and pORCA3-nnLuz (**e**) in transgenic *Nicotiana benthamiana*. The red line is the median, the coloured points represent individual data points. The difference between mean values and *p*-values of post-hoc two-sided Mann–Whitney U tests are indicated above the brackets between the box plots. N = 2-8 biologically independent samples.

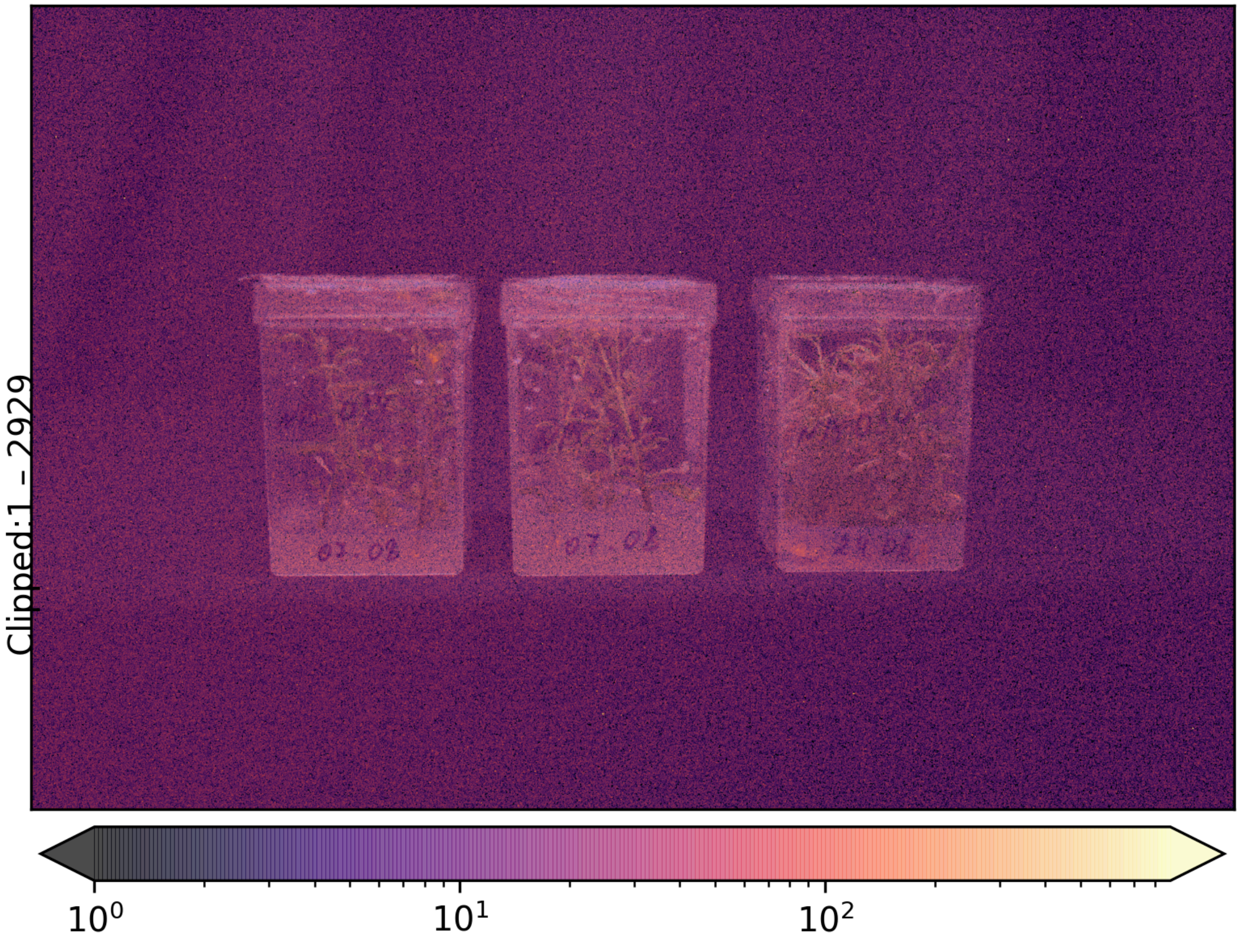


Supplementary Figure 2. Luminescence emitted by the hispidin synthase-lacking *Nicotiana benthamiana* NB034 in vitro plants.
a. Luminescence image shot with Sony alpha, ISO 20000, shutter 30 sec. b. Corresponding to a. image in ambient light. Arrows point to a glowing seedling.

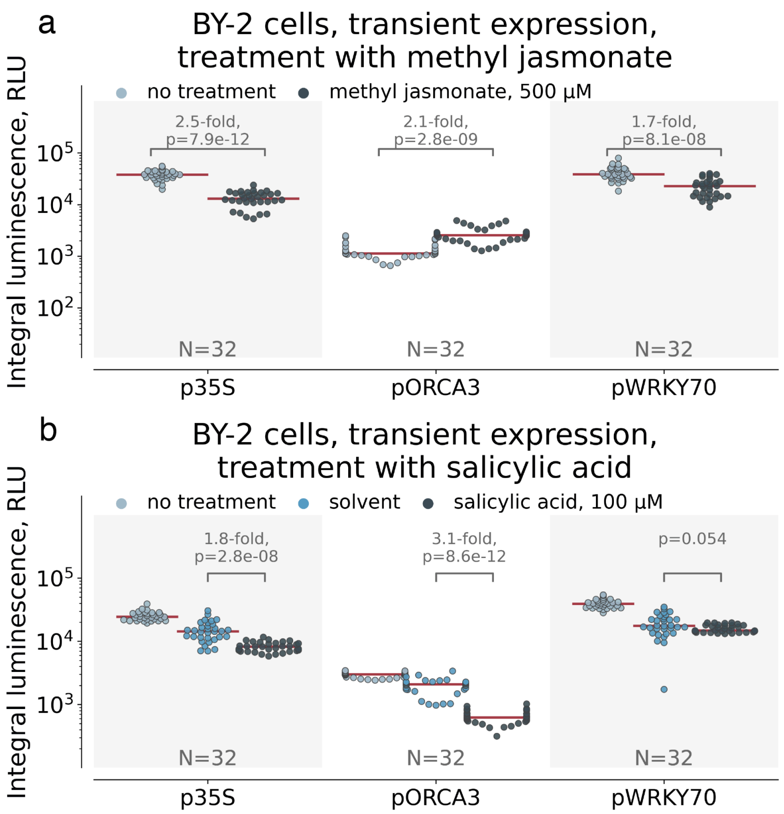


Supplementary Figure 3. Activation of jasmonic and salicylic acid reporters in BY-2.
**a**. Treatment of pORCA3-, pWRKY70- and p35S-nnLuz_v4-expressing BY-2 cells with volatile methyl jasmonate. The red line is the median, the coloured points represent individual data points. The difference between mean values and *p*-values of post-hoc two-sided Mann–Whitney U tests are indicated above the brackets between the box plots. N = 32 plant cell packs.
**b**. Treatment of pORCA3-, pWRKY70- and p35S-nnLuz_v4-expressing BY-2 cells with 100 μM salicylic acid solution. The red line is the median, the coloured points represent individual data points. The difference between mean values and *p*-values of post-hoc two-sided Mann–Whitney U tests are indicated above the brackets between the box plots. N = 32 plant cell packs.

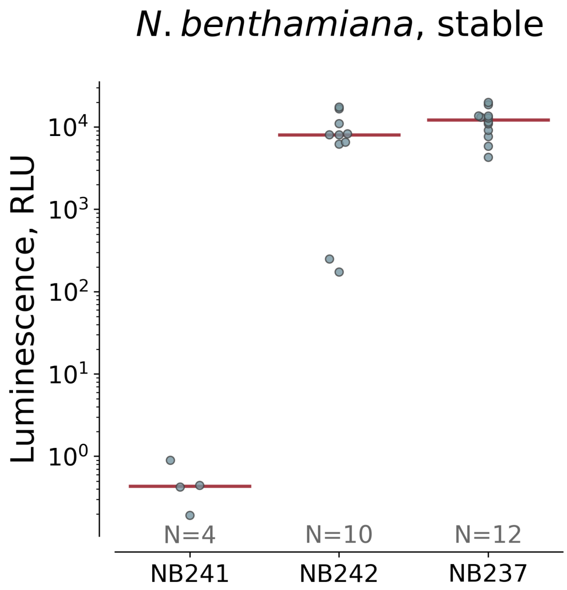


Supplementary Figure 4. Luminescence upon infiltration of leaves of independent luciferase-less *Nicotiana benthamiana* lines with agrobacteria encoding p35S-nnLuz_v4. NB237 was chosen as the masterline for development of other lines used in the project. The red line is the median, the coloured points represent individual data points. N = 4-12 plant cell packs.

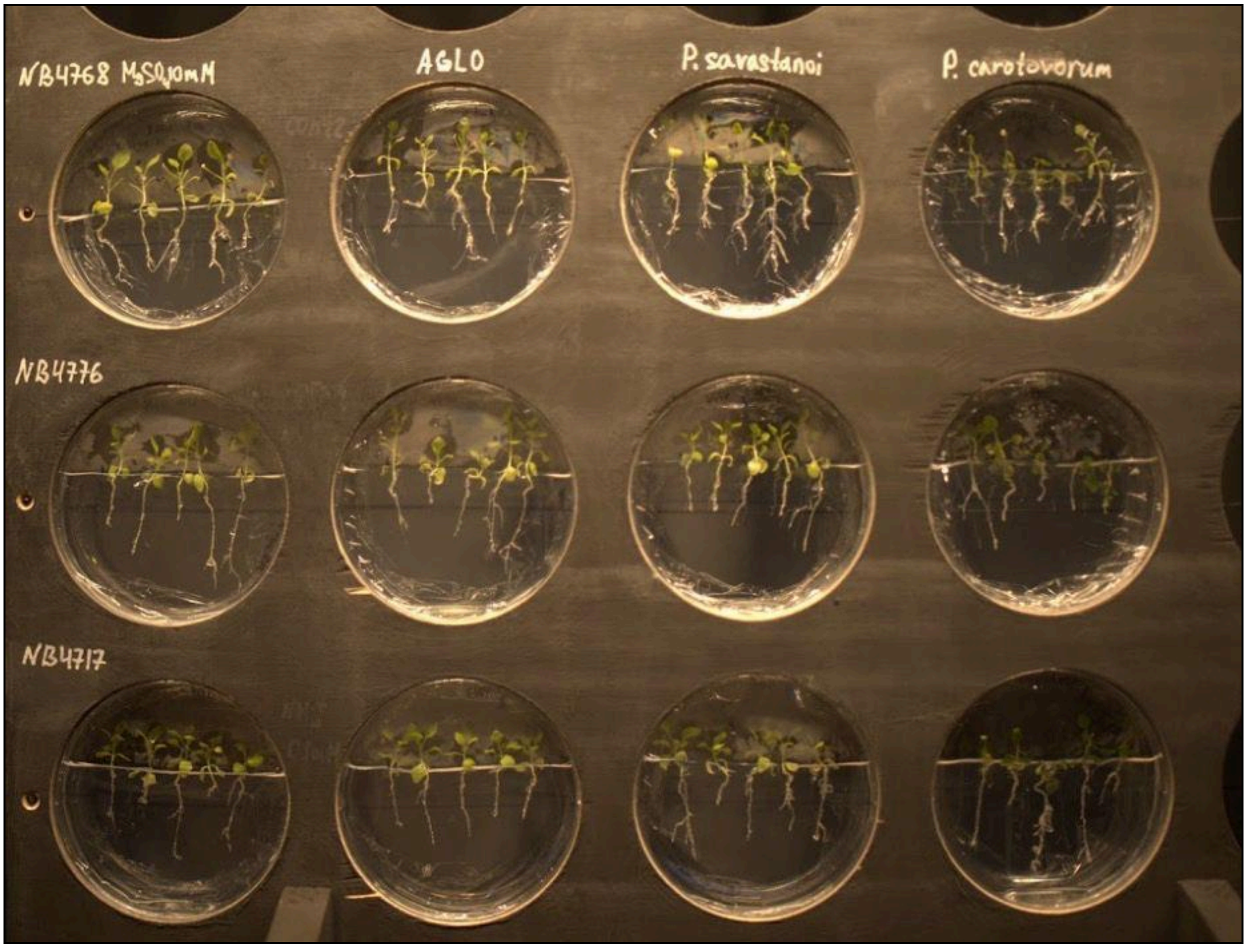


Supplementary Figure 5. A custom wooden setup used to image pathogen infection in plant seedlings.

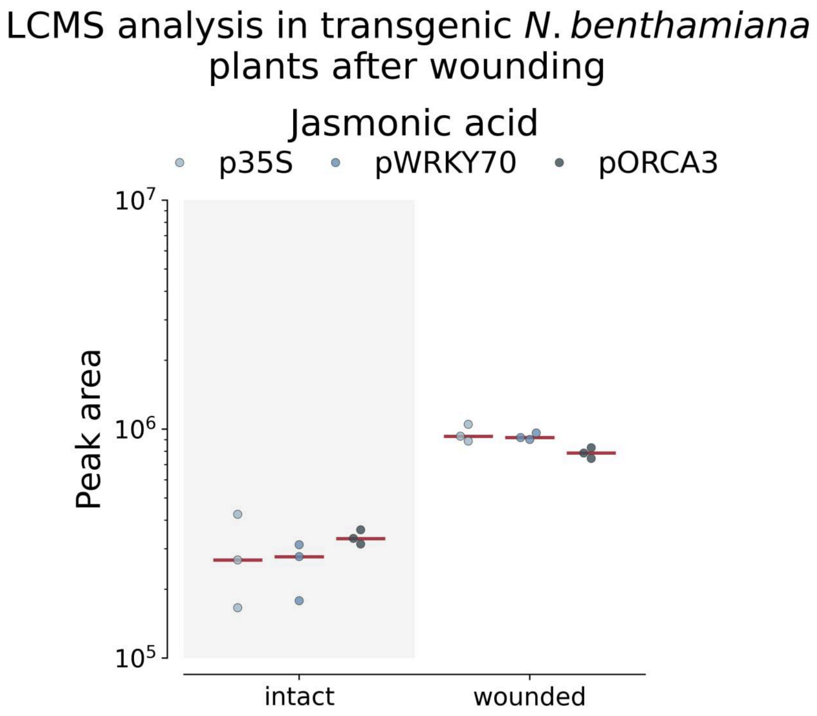


Supplementary Figure 6. The level of jasmonic acid 5 hours after wounding of leaves with scissors, identified with LC-MS. Swarms correspond to biological replicates. The red line is the median, the coloured points represent individual data points. N = 3 biologically independent samples.

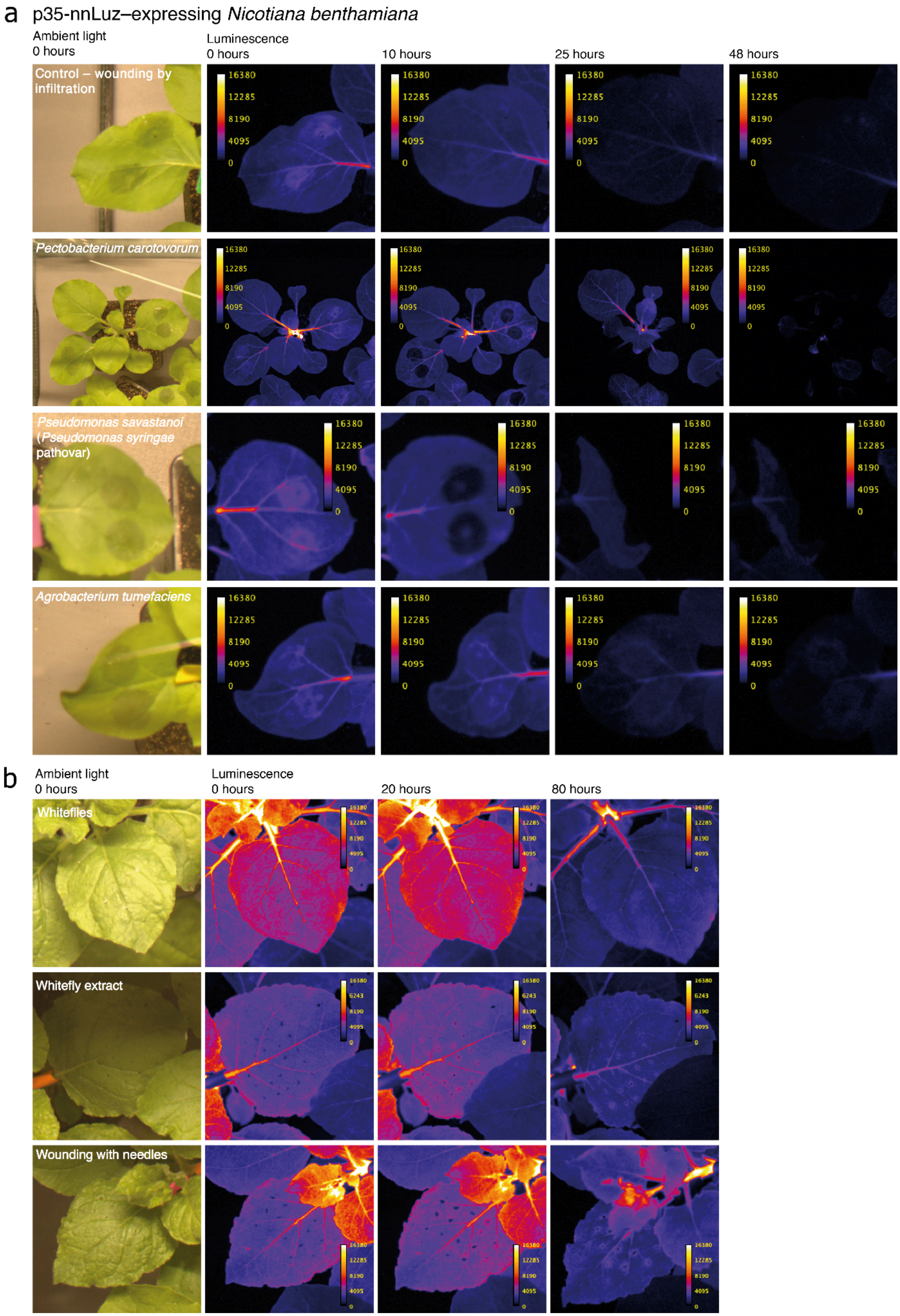

a
p35-nnLuz–expressing *Nicotiana benthamiana*
Ambient light
0 hours
Luminescence
0 hours
10 hours
25 hours
48 hours
Control – wounding by infiltration
*Pectobacterium carotovorum*
*Pseudomonas savastanoi* (*Pseudomonas syringae* pathovar)
*Agrobacterium tumefaciens*
16380
12285
8190
4095
0
b
Ambient light
0 hours
Luminescence
0 hours
20 hours
80 hours
Whiteflies
Whitefly extract
Wounding with needles

Supplementary Figure 7. Luminescence of control p35S-nnLuz-expressing *Nicotiana benthamiana* upon exposure to pests and pathogens.
Luminescence from plants infiltrated with (**a**) buffer, hemibiotroph *Pseudomonas savastanoi*, necrotrophic bacteria *Pectobacterium carotovorum,* or *Agrobacterium tumefaciens.* (**b**) Visible light and luminescence images at different time

points upon exposure to whitefly bites, and to wounding with needles dipped in whitefly extract, or in buffer.

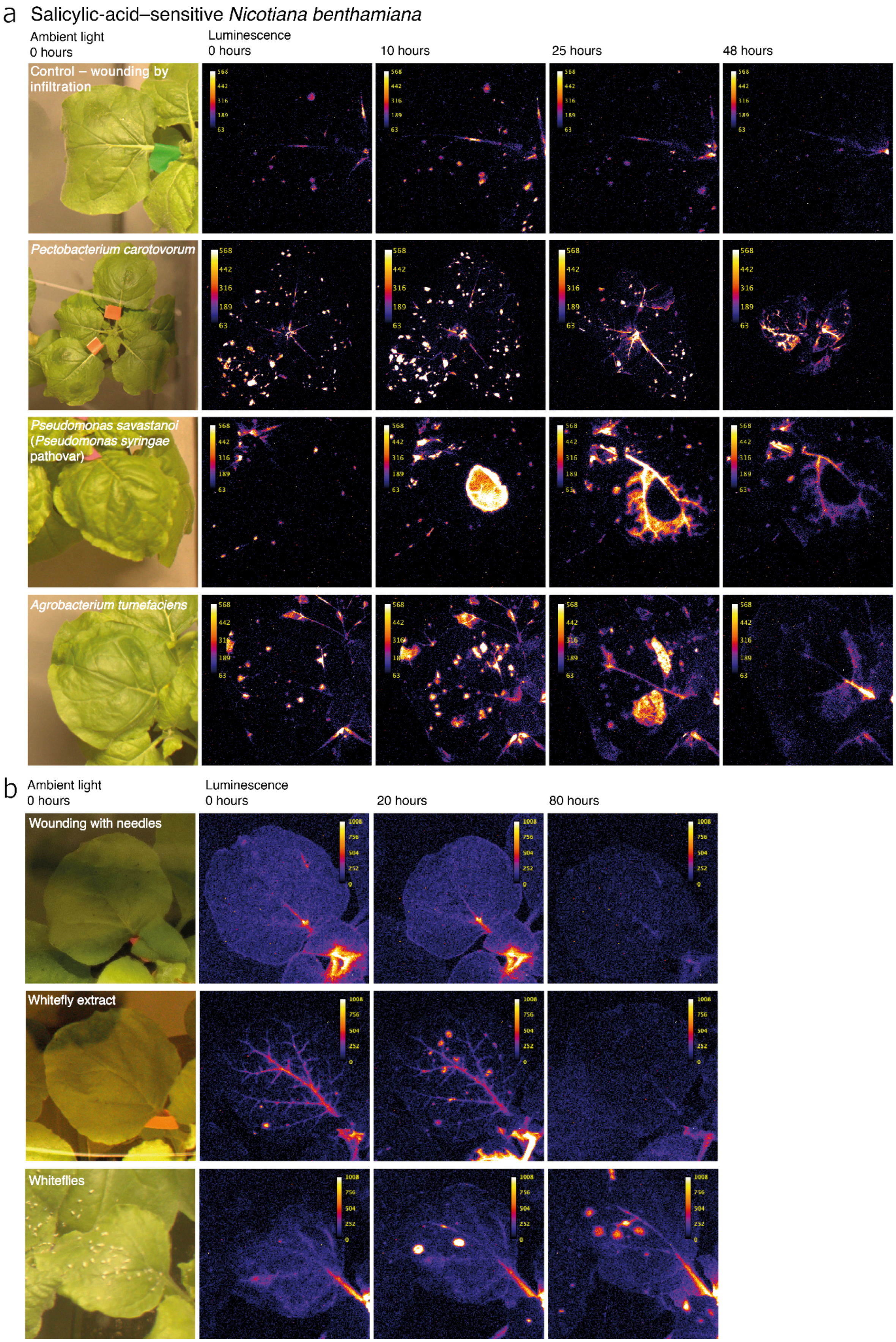

a Salicylic-acid–sensitive Nicotiana benthamiana
Ambient light
0 hours
Luminescence
0 hours
10 hours
25 hours
48 hours
Control – wounding by infiltration
Pectobacterium carotovorum
Pseudomonas savastanoi (Pseudomonas syringae pathovar)
Agrobacterium tumefaciens
568
442
316
189
63
b
Ambient light
0 hours
Luminescence
0 hours
20 hours
80 hours
Wounding with needles
Whitefly extract
Whiteflies
1008
756
504
252
0

Supplementary Figure 8. The response of pWRKY70-nnLuz-expressing transgenic *Nicotiana benthamiana* plants. Visible light image and luminescent images corresponding to different time points for the development of luminescent signals in response to (**a**) infiltration with buffer, hemibiotroph *Pseudomonas savastanoi*, necrotrophic bacteria *Pectobacterium carotovorum,* or *Agrobacterium tumefaciens,* (**b**) whitefly bites, whitefly extract and wounding, or wounding only.

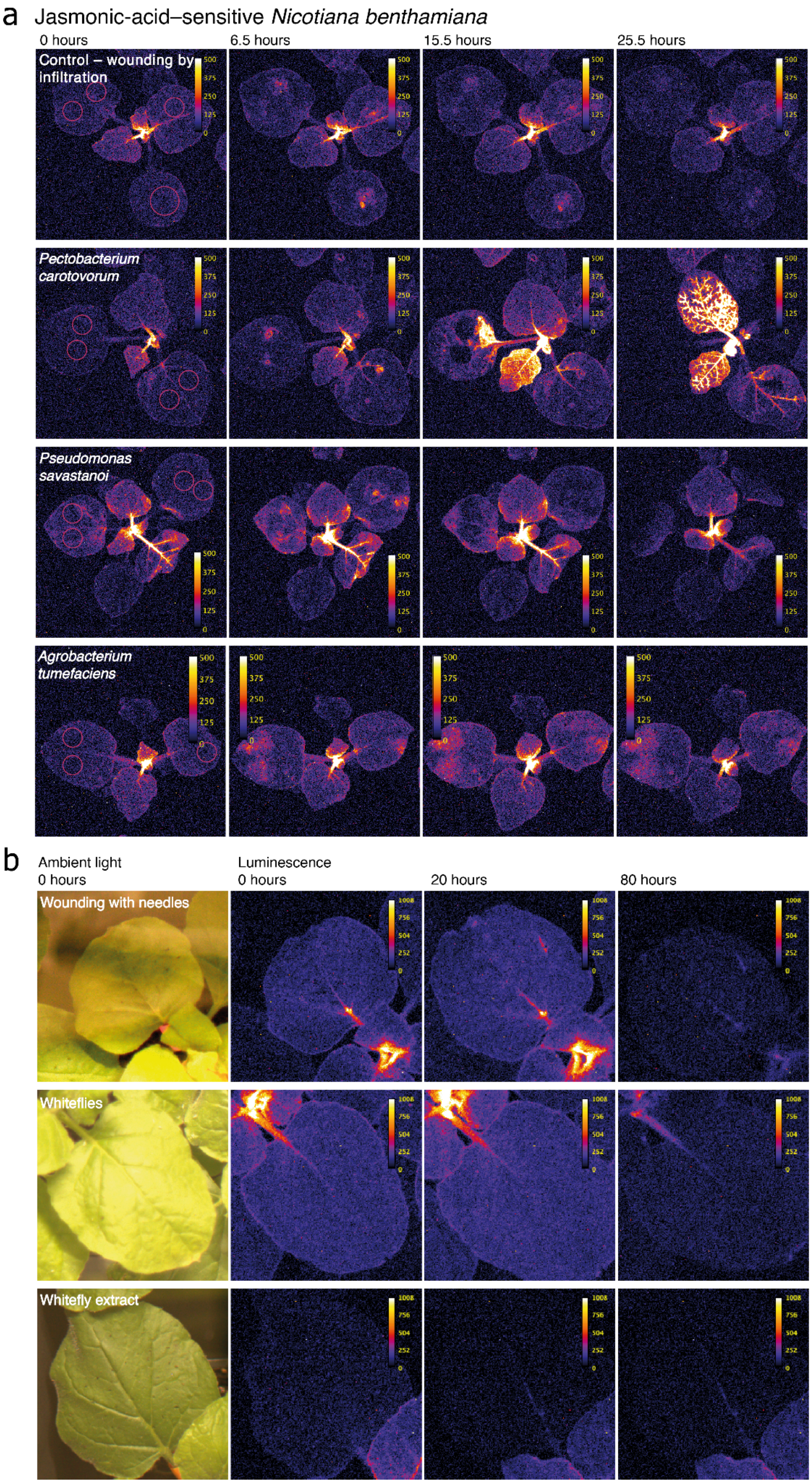
a
Jasmonic-acid–sensitive Nicotiana benthamiana
0 hours
6.5 hours
15.5 hours
25.5 hours
Control – wounding by infiltration
Pectobacterium carotovorum
Pseudomonas savastanoi
Agrobacterium tumefaciens
500
375
250
125
0
b
Ambient light
0 hours
Luminescence
0 hours
20 hours
80 hours
Wounding with needles
Whiteflies
Whitefly extract
1008
756
504
252
0

Supplementary Figure 9. The response of pORCA3-nnLuz-expressing transgenic *Nicotiana benthamiana* plants. Luminescent images corresponding to different time points for the development of luminescent signals in response to (a) infiltration with buffer, hemibiotroph *Pseudomonas savastanoi*, necrotrophic bacteria *Pectobacterium carotovorum,* or *Agrobacterium tumefaciens,* (b) whitefly bites, whitefly extract and wounding, or and wounding only.

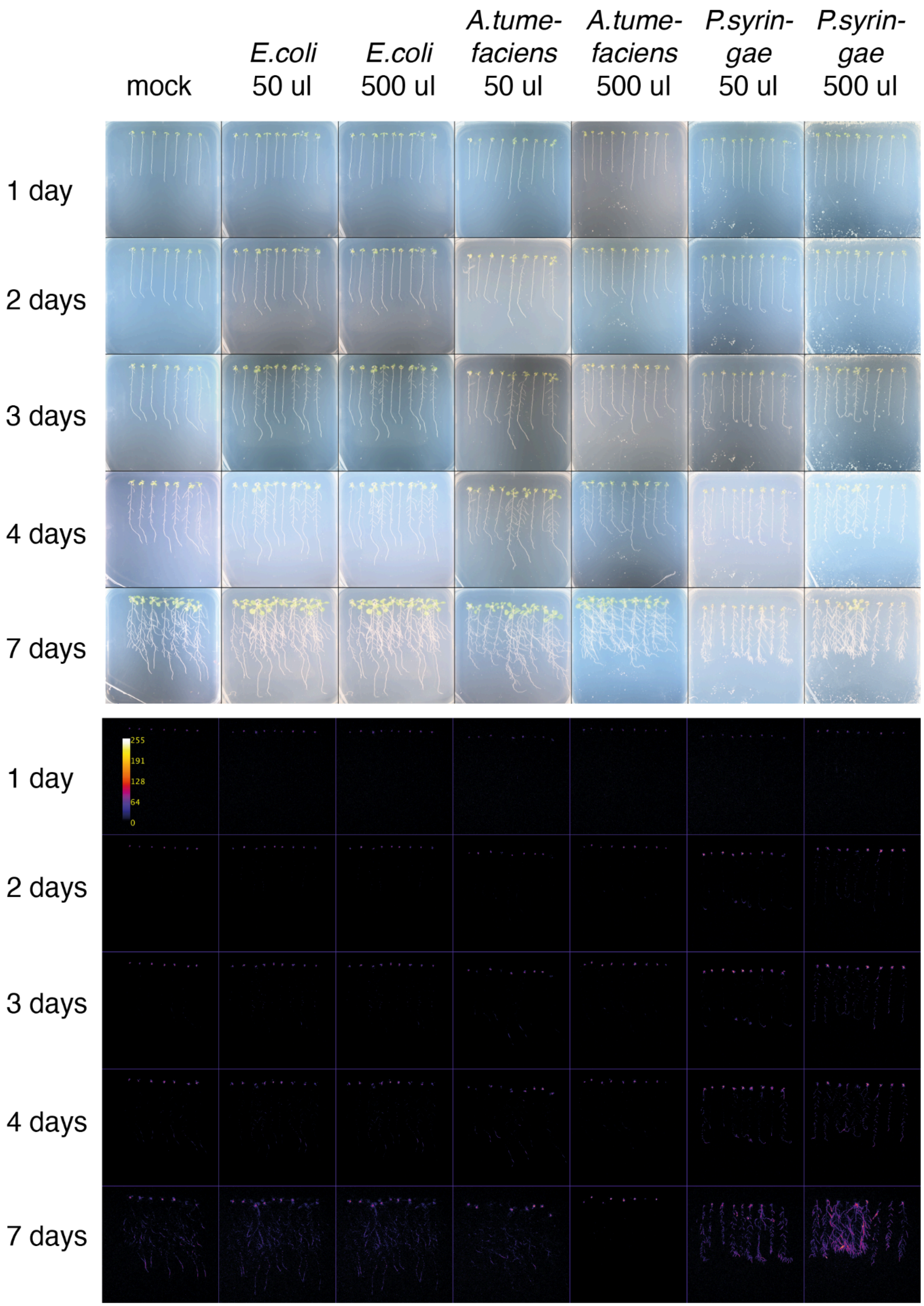
mock
E.coli 50 ul
E.coli 500 ul
A.tume-faciens 50 ul
A.tume-faciens 500 ul
P.syrin-gae 50 ul
P.syrin-gae 500 ul
1 day
2 days
3 days
4 days
7 days
1 day
2 days
3 days
4 days
7 days
255
191
128
64
0

Supplementary Figure 10. The complete imaging of the in vitro infection assay of stably transformed pORCA3 bioluminescence reporters with *E.coli*, *A.tumefaciens*, *P. syringae* DC3000 for 7 days.

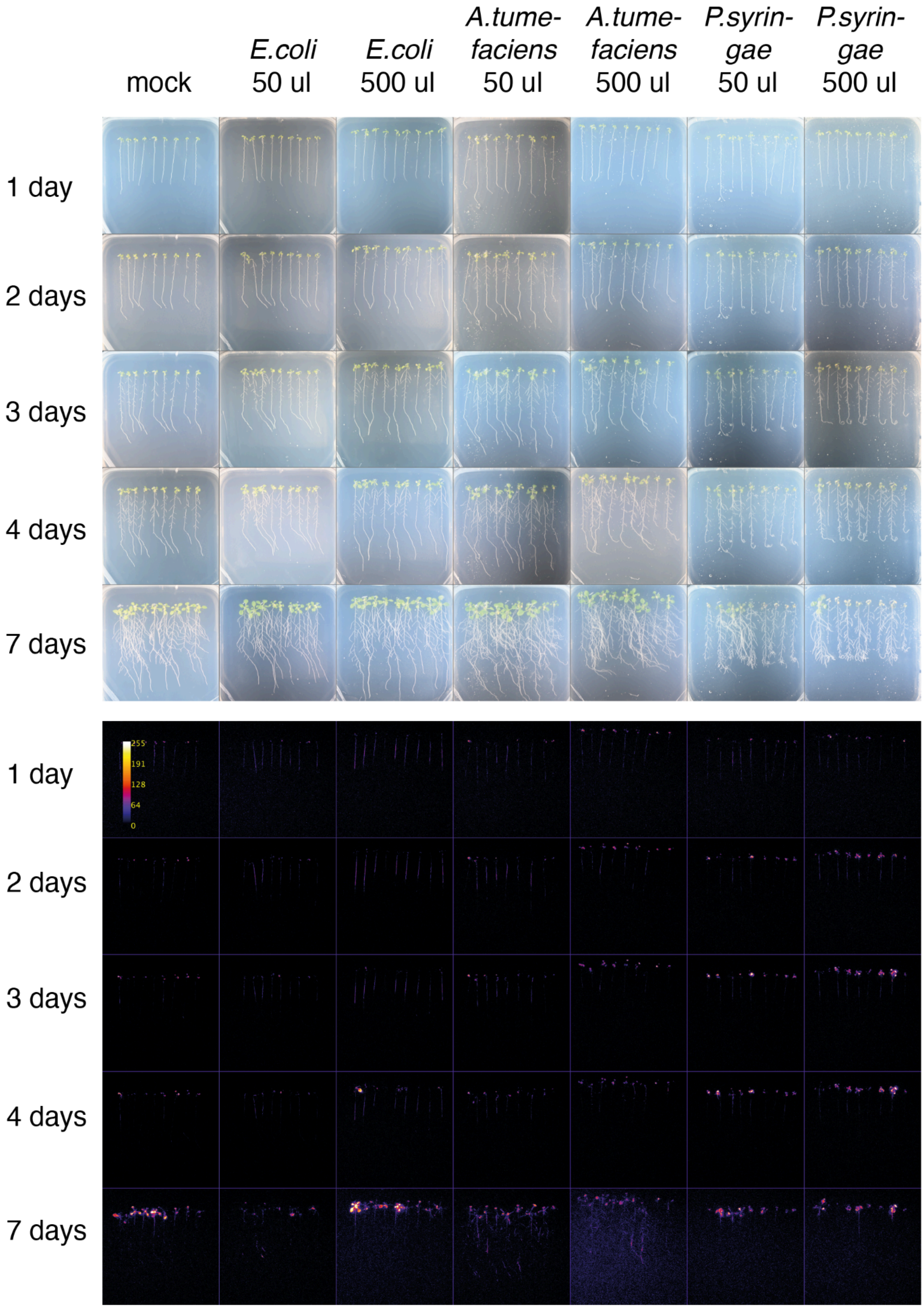
mock
*E.coli* 50 ul
*E.coli* 500 ul
*A.tume-faciens* 50 ul
*A.tume-faciens* 500 ul
*P.syrin-gae* 50 ul
*P.syrin-gae* 500 ul
1 day
2 days
3 days
4 days
7 days
1 day
2 days
3 days
4 days
7 days
255
191
128
64
0

Supplementary Figure 11. The complete imaging of the in vitro infection assay of stably transformed pWRKY70 bioluminescence reporters with *E.coli*, *A.tumefaciens*, *P. syringae* DC3000 for 7 days.

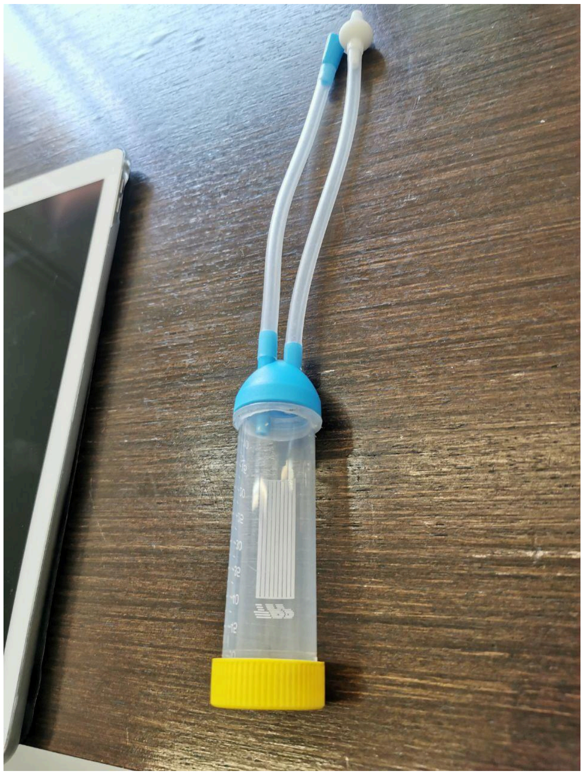

Supplementary Figure 12. A custom aspirator.

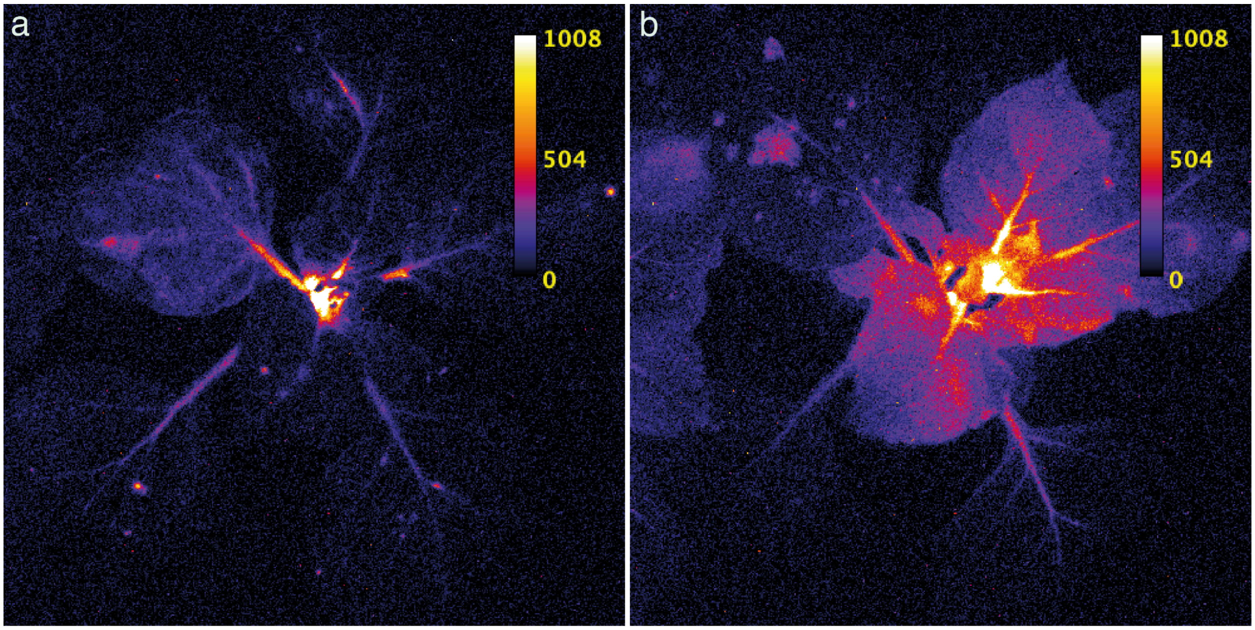


Supplementary Figure 13. Luminescence of salicylic-acid-sensitive plants at the start (a) and at the end (7 day, b) of the whitefly exposure experiment.

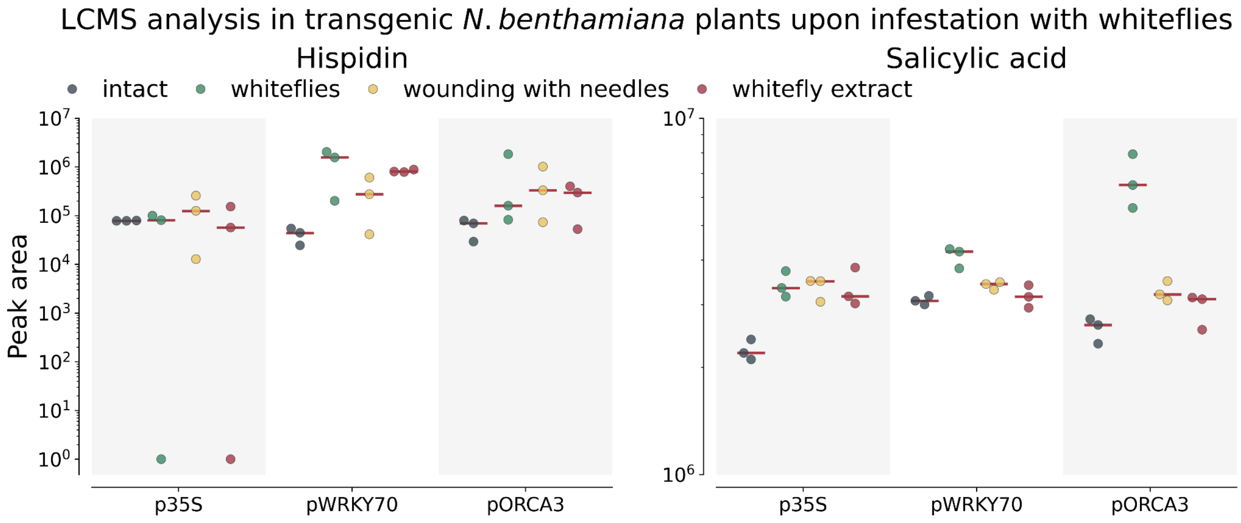


Supplementary Figure 14. The levels of a major fungal bioluminescence metabolite hispidin and salicylic acid upon infestation with whiteflies with LC-MS. The red line is the median, the coloured points represent individual data points.. N = 3 biologically independent samples.

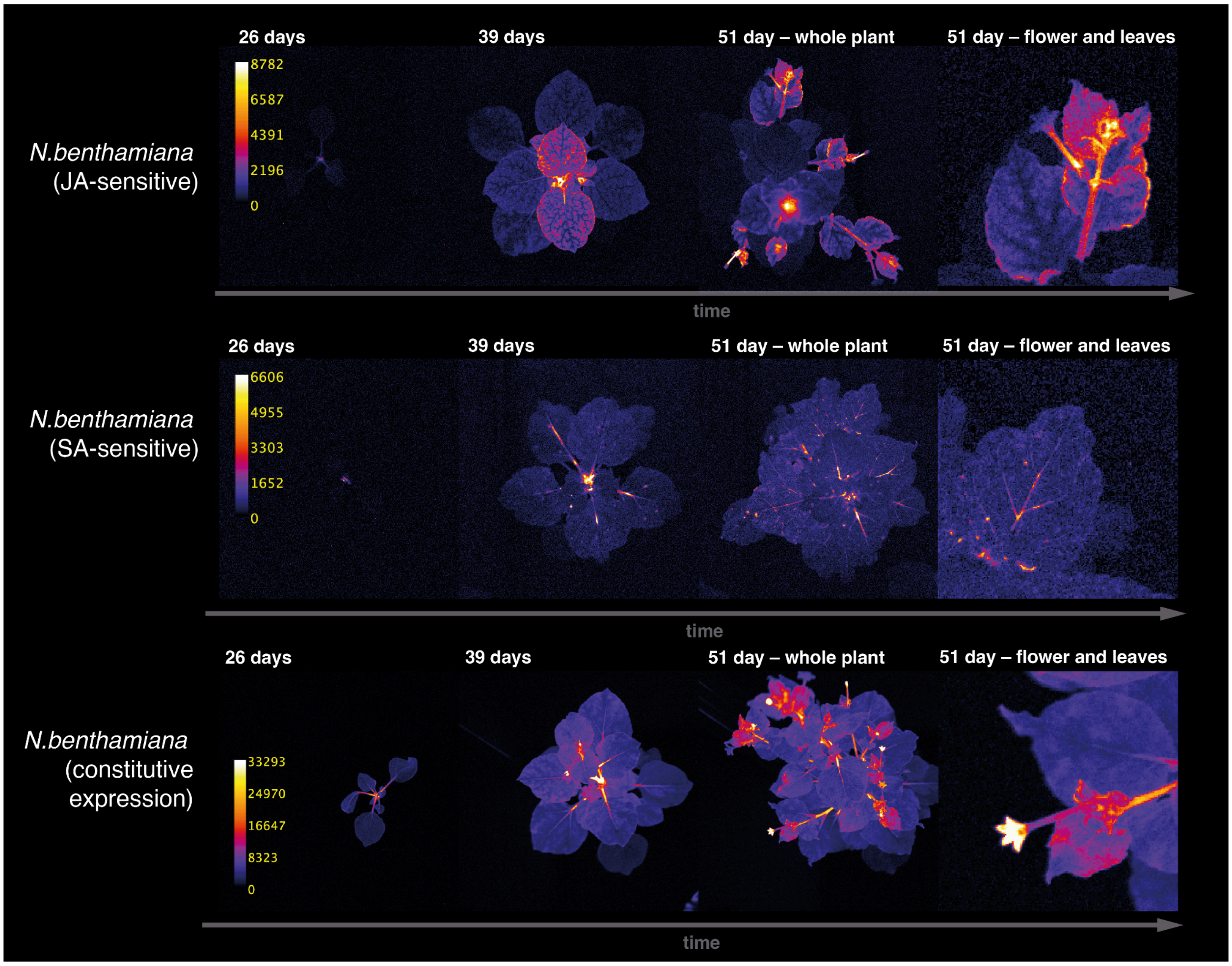


Supplementary Figure 15. Performance of reporters of salicylic, jasmonic acids or constitutively expressing luminescence genes in transgenic *Nicotiana benthamiana* plants during normal growth and flowering.

#### Supplementary Videos

Supplementary Video 1. A 2-day-long timelapse of a pORCA3-nnLuz-expressing *Nicotiana benthamiana* leaf upon wounding – infiltration with buffer (upper left), infiltration of *Pseudomonas savastanoi* (upper right), infiltration of *Pectobacterium carotovorum* (lower left), and infiltration of *Agrobacterium tumefaciens* (lower right). Shot with Sony Alpha ILCE-7M3 camera and 35-mm T1.5 ED AS UMC VDSLR lens (Samyang, ~f/1.4) with an exposure of 30 s and ISO 3,200. Imaging performed every 30 minutes.

Supplementary Video 2. A 2-day-long timelapse of a p35S-nnLuz-expressing *Nicotiana benthamiana* leaf upon wounding – infiltration with buffer (upper left), infiltration of *Pseudomonas savastanoi* (upper right), infiltration of *Pectobacterium carotovorum* (lower left), and infiltration of *Agrobacterium tumefaciens* (lower right). Shot with Sony Alpha ILCE-7M3 camera and 35-mm T1.5 ED AS UMC VDSLR lens (Samyang, ~f/1.4) with an exposure of 30 s and ISO 3,200. Imaging performed every 30 minutes.

Supplementary Video 3. A 2-day-long timelapse of a pWRKY70-nnLuz-expressing *Nicotiana benthamiana* leaf upon wounding – infiltration with buffer (upper left), infiltration of *Pseudomonas savastanoi* (upper right), infiltration of *Pectobacterium carotovorum* (lower left), and infiltration of *Agrobacterium tumefaciens* (lower right). Shot with Sony Alpha ILCE-7M3 camera and 35-mm T1.5 ED AS UMC VDSLR lens (Samyang, ~f/1.4) with an exposure of 30 s and ISO 3,200. Imaging performed every 30 minutes.

Supplementary Video 4. A 7-day-long timelapse of a p35S-nnLuz-expressing *Nicotiana benthamiana* leaf after wounding with needles. Shot with Sony Alpha ILCE-7M3 camera and 35-mm T1.5 ED AS UMC VDSLR lens (Samyang, ~f/1.4) with an exposure of 30 s and ISO 3,200. Imaging performed every 30 minutes.

Supplementary Video 5. A 7-day-long timelapse of a p35S-nnLuz-expressing *Nicotiana benthamiana* leaf after wounding with needles and treatment with whitefly extract. Shot with Sony Alpha ILCE-7M3 camera and 35-mm T1.5 ED AS UMC VDSLR lens (Samyang, ~f/1.4) with an exposure of 30 s and ISO 3,200. Imaging performed every 30 minutes.

Supplementary Video 6. A 7-day-long timelapse of a p35S-nnLuz-expressing *Nicotiana benthamiana* leaf after whitefly infestation. Shot with Sony Alpha ILCE-7M3 camera and 35-mm T1.5 ED AS UMC VDSLR lens (Samyang, ~f/1.4) with an exposure of 30 s and ISO 3,200. Imaging performed every 30 minutes.

Supplementary Video 7. A 7-day-long timelapse of a pORCA3-nnLuz-expressing *Nicotiana benthamiana* leaf after wounding with needles. Shot with Sony Alpha ILCE-7M3 camera and 35-mm T1.5 ED AS UMC VDSLR lens (Samyang, ~f/1.4) with an exposure of 30 s and ISO 3,200. Imaging performed every 30 minutes.

Supplementary Video 8. A 7-day-long timelapse of a pORCA3-nnLuz-expressing *Nicotiana benthamiana* leaf after wounding with needles and treatment with whitefly extract. Shot with Sony Alpha ILCE-7M3 camera and 35-mm T1.5 ED AS UMC VDSLR lens (Samyang, ~f/1.4) with an exposure of 30 s and ISO 3,200. Imaging performed every 30 minutes.

Supplementary Video 9. A 7-day-long timelapse of a pORCA3-nnLuz-expressing *Nicotiana benthamiana* leaf after whitefly infestation. Shot with Sony Alpha ILCE-7M3 camera and 35-mm T1.5 ED AS UMC VDSLR lens (Samyang, ~f/1.4) with an exposure of 30 s and ISO 3,200. Imaging performed every 30 minutes.

Supplementary Video 10. A 7-day-long timelapse of a pWRKY70-nnLuz-expressing *Nicotiana benthamiana* leaf after wounding with needles. Shot with Sony Alpha ILCE-7M3 camera and 35-mm T1.5 ED AS UMC VDSLR lens (Samyang, ~f/1.4) with an exposure of 30 s and ISO 3,200. Imaging performed every 30 minutes.

Supplementary Video 11. A 7-day-long timelapse of a pWRKY70-nnLuz-expressing *Nicotiana benthamiana* leaf after wounding with needles and treatment with whitefly extract. Shot with Sony Alpha ILCE-7M3 camera and 35-mm T1.5 ED AS UMC VDSLR lens (Samyang, ~f/1.4) with an exposure of 30 s and ISO 3,200. Imaging performed every 30 minutes.

Supplementary Video 12. A 7-day-long timelapse of a pWRKY70-nnLuz-expressing *Nicotiana benthamiana* leaf after whitefly infestation. Shot with Sony Alpha ILCE-7M3 camera and 35-mm T1.5 ED AS UMC VDSLR lens (Samyang, ~f/1.4) with an exposure of 30 s and ISO 3,200. Imaging performed every 30 minutes.

Supplementary Video 13. A 1.5-month-long timelapse of a p35S-nnLuz-expressing *Nicotiana benthamiana* plant. Shot with Sony Alpha ILCE-7M3 camera and 35-mm T1.5 ED AS UMC VDSLR lens (Samyang, ~f/1.4) with an exposure of 30 s and ISO 400. Imaging performed every 30 minutes.

Supplementary Video 14. A 1.5-month-long timelapse of a pORCA3-nnLuz-expressing *Nicotiana benthamiana* plant. Shot with Sony Alpha ILCE-7M3 camera and 35-mm T1.5 ED AS UMC VDSLR lens (Samyang, ~f/1.4) with an exposure of 30 s and ISO 400. Imaging performed every 30 minutes.

Supplementary Video 15. A 1.5-month timelapse of a pWRKY70-nnLuz-expressing *Nicotiana benthamiana* plant. Shot with Sony Alpha ILCE-7M3 camera and 35-mm T1.5 ED AS UMC VDSLR lens (Samyang, ~f/1.4) with an exposure of 30 s and ISO 400. Imaging performed every 30 minutes.

Supplementary Table 1. Plasmids used in the study.

| Plasmid Name | Purpose | Gene or insert Name | Sequence |
|---|---|---|---|
| pNK091 | MoClo-compatible Level P vector carrying fungal bioluminescence system lacking luciferase gene | pNos-KanR-ocsT \| p35S-nnHispS-ocsT \| pCmYLCV-npgA-ATPT \| p35S-nnCPH- ocsT \| pFMV-nnH3H-nosT | https://benchling.com/s/seq-JChGOcCVNiYRKbruZQ2z?m=slm-cN2aKkBzOeeBNZLMgSwe |
| pNK3625 | MoClo-compatible Level 0 vector carrying promoter pWRKY70 | Promoter WRKY70 | https://benchling.com/s/seq-LnX4S8GvV3xHMzRjPqmV?m=slm-66eAnYZagmCqJt7dTydy |
| M6597 | MoClo-compatible Level 0 vector carrying terminator WRKY70 | Terminator WRKY70 | https://benchling.com/s/seq-zClxpyzVT87rb6uvtRUp?m=slm-lVqcIu56Z2ITnbpnnJ7q |
| M6667 | MoClo-compatible Level 0 vector carrying terminator ORCA3 | Terminator ORCA3 | https://benchling.com/s/seq-uqqsyEjt3uX8uHkYEMEB?m=slm-sNaSY3v0tTCZrDSmSeMK |
| M6570 | MoClo-compatible Level 0 vector carrying promoter pORCA3 | Promoter ORCA3 | https://benchling.com/s/seq-MXTjKQCLrWJmQlmLTNL9?m=slm-e9BQnrrx5LKiSVoaebK2 |
| M7138 | MoClo-compatible Level M vector encoding hygromycin resistance cassette and a luciferase gene under control of ORCA3 promoter and terminator | pNOS-HygR-ocsT \| pORCA3-nnLuz-ORCA3_T | https://benchling.com/s/seq-7VQRjD9A4fue1L0KthDX?m=slm-V97liy3kx0KRKInU8HLU |
| M7691 | MoClo-compatible Level M vector encoding hygromycin resistance cassette and a luciferase gene under control of WRKY70 promoter and | pNOS-HygR-ocsT \| pWRKY70-nnLuz-WRKY70_T | https://benchling.com/s/seq-g4ASG5qVvLKQFVUu2Qyw?m=slm-3AaEmS8nTghh4GK05rlm |

| | | | |
|---|---|---|---|
| pNK3558 | MoClo-compatible Level M vector encoding hygromycin resistance cassette and a luciferase gene under control of p35S promoter and act2 terminator | pNOS-HygR-ocsT \| p35S-nnLuz-act2 | https://benchling.com/s/seq-YmmkwEubHTyo7FL1nbtv?m=slm-DjOwagVTlD5X6nYe0PdN |

**Supplementary Table 2.** Plant lines used in the study.

| Line ID | Line name | Inserts |
|---|---|---|
| NB237 | *N. benthamiana* luciferase-less masterline | pNos-KanR-ocsT \| p35S-nnHispS-ocsT \| pCmYLCV-npgA-ATPT \| p35S-nnCPH-ocsT \| pFMV-nnH3H-nosT |
| NB4717 | *N. benthamiana* salicylic-acid-sensitive line | pNos-KanR-ocsT \| p35S-nnHispS-ocsT \| pCmYLCV-npgA-ATPT \| p35S-nnCPH-ocsT \| pFMV-nnH3H-nosT<br><br>pNOS-HygR-ocsT \| pWRKY70-nnLuz-WRKY70_T |
| NB4776 | *N. benthamiana* jasmonic-acid-sensitive line | pNos-KanR-ocsT \| p35S-nnHispS-ocsT \| pCmYLCV-npgA-ATPT \| p35S-nnCPH-ocsT \| pFMV-nnH3H-nosT<br><br>pNOS-HygR-ocsT \| pORCA3-nnLuz-ORCA3_T |
| NB4768 | *N. benthamiana* autoluminescent line | pNos-KanR-ocsT \| p35S-nnHispS-ocsT \| pCmYLCV-npgA-ATPT \| p35S-nnCPH-ocsT \| pFMV-nnH3H-nosT<br><br>pNOS-HygR-ocsT \| p35S-nnLuz-act2T |
| AT8462 | *A. thaliana* luciferase-less masterline | pNos-KanR-ocsT \| p35S-nnHispS-ocsT \| pCmYLCV-npgA-ATPT \| p35S-nnCPH-ocsT \| pFMV-nnH3H-nosT |
| AT8463 | *A. thaliana* salicylic-acid-sensitive line | pNos-KanR-ocsT \| p35S-nnHispS-ocsT \| pCmYLCV-npgA-ATPT \| p35S-nnCPH-ocsT \| pFMV-nnH3H-nosT<br><br>pNOS-HygR-ocsT \| pWRKY70-nnLuz-WRKY70_T |
| AT8464 | *A. thaliana* jasmonic-acid-sensitive line | pNos-KanR-ocsT \| p35S-nnHispS-ocsT \| pCmYLCV-npgA-ATPT \| p35S-nnCPH-ocsT \| pFMV-nnH3H-nosT<br><br>pNOS-HygR-ocsT \| pORCA3-nnLuz-ORCA3_T |
